# Ventral anterior foregut endoderm cells as progenitors for airway epithelial cell replacement in primary ciliary dyskinesia

**DOI:** 10.64898/2026.02.05.701025

**Authors:** Carine Bourdais, Agathe Cœur, Florent Foisset, Marion Nadaud, Cecilia Urena, Amel Nasri, Joffrey Mianné, Lisa Morichon, Fanny Rolland, Lounis Yakhou, Aurélie Petit, Qiang Bai, Isabelle Vachier, Said Assou, Arnaud Bourdin, John De Vos

## Abstract

**Background:** Lung transplantation remains the ultimate treatment option for patients with end-stage lung disease, but has many limitations. This underlines the urgent need of developing alternative approaches to treat lung disorders. Among the emergent strategies, gene therapy holds great potential for the treatment of monogenic lung diseases. However, so far, aerosolized viral vector-based delivery for gene therapy has failed likely because of difficulties in accessing the target cells. Combined gene and cell therapy approaches could be a promising alternative. Trials using basal cell transplantation already showed encouraging results. Moreover, the induced pluripotent stem cell (iPSC) technology broadens the scope of personalized therapies by paving the way for autologous approaches. Our group previously derived iPSC lines from patients with Primary Ciliary Dyskinesia (PCD) and found that their correction by gene conversion allows functional recovery. This study aimed to identify the best progenitors and airway conditioning technique to develop an autologous cell replacement strategy for PCD.

**Methods:** Airway epithelial cells were differentiated from induced pluripotent stem cell (iPSC) lines from a healthy donor (parental Hy03) and Hy03 in which *MCIDAS* was knocked out (PCD model) and maintained in air-liquid interface (iALI). The engraftment of GFP+ ventral Anterior Foregut Endoderm (vAFE) cells, differentiated from GFP-expressing Hy03 iPSCs, was assessed after conditioning of the recipient iALI. The efficacy (epithelial cell shedding) and toxicity (cell death) of different conditioning strategies were compared. Cilia functional repair was assessed using microbead motion tracking.

**Results:** GFP+ vAFE cells can successfully integrate and repair trypsin- or EDTA-conditioned airway epithelia derived from the parental and *MCIDAS*^-/-^ Hy03 iPSC lines. EDTA showed optimal efficacy/safety balance. Progenitor integration and differentiation were confirmed by E-cadherin, tubulin-IV, KRT5 and MUC5AC co-expression in GFP+ engrafted cells at day 35 post-graft (immunofluorescence analysis). The engrafted GFP+ population reached 35-45% of the total epithelial population, as indicated by flow cytometry quantification of EpCAM+/GFP+ cells. Functional analysis demonstrated cilia motion restoration after GFP+ cell engraftment onto *MCIDAS*^*-/-*^ iALI.

**Conclusions:** Our study shows that vAFE cells can integrate and differentiate to repair epithelial models of PCD. EDTA conditioning is promising for the clinical application of this therapeutic strategy.

## Background

Primary Ciliary Dyskinesia (PCD) is a rare monogenic lung condition in which genetic mutations impair cilia function (1–3). PCD estimated incidence is ∼1 in 10 000 live births. Currently, disease-causing variants have been identified in ∼50 genes (4,5). In the most severe cases, lung transplantation can be required (6). For instance, pathogenic mutations in the *MCIDAS* gene are among the most deleterious because of the key role played by multicilin (the protein encoded by this gene) in cilia generation (7).

Several studies demonstrated that it is possible to restore cilia activity using gene therapy (8,9). However, multiple challenges prevent its immediate clinical application, including limited transgene integration and the potential immune response (10,11). As treatment of genetic lung diseases using aerosolized viral vector-based gene therapy has failed, combined gene and cell therapy has emerged as an alternative (12,13).

In cell-based therapies, the patient’s cells are corrected *ex vivo* by gene editing before replacement. The safety of the engineered cells can be assessed before transplantation, and the use of autologous cells prevents immune rejection. Currently, the main challenges of this approach are the identification of the most suitable cells for achieving and maintaining long-term repair and the development of strategies to optimize their integration (12).

In lung diseases, long-term repair implies replacing progenitor cells. KRT5+TP63+ adult basal cells can self-renew and differentiate, including into ciliated cells (14–17). They can be isolated from patient biopsies, but their *ex vivo* expansion after gene editing appears challenging for clinical applications due to their heterogeneity in terms of epithelial repair capacity (18–20). Alternatively, induced pluripotent stem cells (iPSC) offer a new and infinite cell source for autologous cell therapy (21–23). After reprogramming, iPSCs can be quite easily engineered to correct the genetic alteration (24,25). Then, by mimicking lung embryonic development, iPSCs can be differentiated into the different cell types of the bronchial epithelium, including basal cells, but also into earlier progenitors such as ventral anterior foregut endoderm (vAFE) cells (26–30). In theory, vAFE cells offer advantages similar to basal cells for therapeutic applications, including a broad differentiation capacity. In addition, they can be produced faster and in higher number and should contain key hillock basal stem cells that facilitate long-lasting cell replacement (31,32). In this proof-of-concept study, we assessed the ability of early lung progenitors (vAFE cells), differentiated from iPSCs, to functionally repair a PCD model.

## Methods

### iPSC lines, culture and passaging

The human iPSC line Hy03 (UHOMi002-A) was generated in our laboratory and maintained in culture as previously reported (33). From this cell line, two models were generated: (1) a cell line in which *MCIDAS* was knocked out (Hy03 MCIDAS^-/-^) to mimic the PCD bronchial epithelium that will receive the cell graft, and (2) a cell line that constitutively express green fluorescent protein (Hy03 GFP) under the control of the EF1a promoter as a model of autologous cell donor (25).

Briefly, the iPSC lines were maintained in undifferentiated stage in feeder-free conditions on Geltrex™ growth factor-reduced matrix (Gibco, A1413202) in Essential 8™ medium (Gibco, A1517001) that was changed daily. Cells were grown at 37°C and 5% CO_2_. Colonies were passaged when they reached 70-90% of confluence (every ∼4 days) using Versene (Gibco, 15040066). Single cells were harvested and plated at a ratio of 1:10 to 1:20 with the ROCK inhibitor Y-27632 (Biotechne, 1254).

### Differentiation of iPSCs into bronchial epithelium

Then, iPSCs were differentiated into functional bronchial epithelium in air-liquid interface (ALI) resulting in iPSC-derived ALI (iALI), as previously described (26). Briefly, cells were plated at 80 000 cells/cm^2^ on Geltrex™-coated culture plates in Essential 8™ medium. In the first 7 days, medium was changed each day. RPMI 1640 (Gibco, 21875034) was supplemented with B-27 (without vitamin A; Gibco, 12587010) and different factors were added to induce pathways involved in the development of the anterior primitive streak, the definitive endoderm. and the vAFE. At day 7, when NKX2.1 expression is peaking, cells were harvested in large clumps using a culture rake and plated in Transwell™ plates (Corning™, 3460) coated with Geltrex™, in Pneumacult™-ExPlus Medium (Stemcell Technologies, 05040) for 2 days, and then in Pneumacult™-ALI Basal Medium (Stemcell Technologies, 05001). Cells were polarized after 2 days in Pneumacult™-ALI Basal Medium, and DAPT (Notch inhibitor; Biotechne, 2634) was added to the medium 2 weeks after polarization to drive cell maturation towards ciliated cells.

The differentiation process was monitored at day 3 (expression of CXCR4 and FOXA2 from cells of the definitive endoderm) and at day 7 (expression of NKX2.1 and SOX2 by vAFE cells) by flow cytometry analysis.

### Bronchial epithelium conditioning and donor cell graft

After 5 weeks of differentiation, mature iALI epithelia were conditioned to allow lung progenitor cell engraftment. After removing the medium, iALI basal and apical surfaces were washed with PBS (Gibco, 20012068). Then, the different conditioning compounds listed in Table S1 were added at the apical side of the model for the indicated time (Table S1) at 37°C and 5% CO_2_. After washing with PBS, Pneumacult™-ALI Basal Medium was added to the basal side. Concomitantly, Hy03 GFP lung progenitor cells (at day 7 of differentiation; donor cells) were harvested with Versene or Trypsin-EDTA 0.25% solution (Gibco, 25200056) in single cells or clumps and suspended in Pneumacult™-ExPlus Medium. Then, they were plated at the desired cell concentration, in a final volume of 200 µL, at the conditioned iALI apical surface. The remaining supernatant was aspirated after 24 h to re-polarize the model, and Pneumacult™-ALI Basal Medium was changed every day at the basal side.

### Paraffin sections

For histological analysis, iALI samples were fixed in a 4% paraformaldehyde solution (Thermo Scientific, 28908) at 4°C for 24 h after washing with PBS solution. After dehydration with increasing concentrations of ethanol, samples were embedded in paraffin using the Myr STP120 automate and the Myr EC350 station (following the manufacturer’s instructions). Then, sections (3 to 8 µm) were cut with a microtome (Leica, RM2125 RTS) and mounted on microscope slides (Epredia, AG00008032E01FST20, J7850AMNZ, or J2800AMNZ).

### Immunofluorescence staining

For immunofluorescence staining, sections were deparaffinized twice in Neo-clear™ solution (Sigma-Aldrich, 1.09843) and ethanol solutions at decreasing concentration and then incubated in a solution of citrate buffer (Sigma-Aldrich, C9999) in a water bath at 100°C for 20 min. Autofluorescence was reduced by incubation with 0.06% potassium permanganate solution at room temperature (RT) for 20 min. Then, cells were permeabilized with 0.5% Triton X-100 at RT for 15 min and blocked with 10% donkey serum (Sigma-Aldrich, S30-M) diluted in 1% BSA/0.1% Triton X-100/PBS solution (staining buffer; Sigma-Aldrich, 17906; Thermo Scientific Chemicals, A16046.AE) at RT for 1h before incubation with the primary antibodies diluted at the appropriated concentration in the staining buffer at 4°C overnight. Antibodies are listed in Table S2. Then, sections were washed with 0.025% Triton X-100/PBS and incubated with the secondary antibodies (1:1000 in staining buffer) at RT in the dark for 1 h. Lastly, nuclei were stained with DAPI (Sigma-Aldrich, D9542; 1:2500 in PBS) at RT, in the dark, for 2 min and sections were mounted in aqueous resin (BIO-RAD, BUF058B) and stored in the dark at 4°C till image acquisition.

Images were acquired with a TCS SP5 or TCS SP8 confocal microscope (Leica), BioTek Cytation 5 reader (Agilent) or Axioscan automate (ZEISS). ImageJ (Fiji) or Gen5 was used, respectively, for analysis.

### iALI dissociation and flow cytometry analysis

To harvest iALI in single cell suspensions, samples were washed with PBS and incubated with pre-warmed 0.25% trypsin solution, with/without dispase (Sigma-Aldrich, D4693) or collagenases (Gibco, 17018029; 17101015; 17104019), at 37°C for 8-10 min. Detached cells were resuspended in 10% foetal bovine serum (diluted in PBS or Versene solution) and incubated with trypsin 2 to 4 times until complete dissociation. Pipetting or filtration helped to fully dissociate the remaining aggregates. Cell suspensions were centrifuged at 300g for 5 min and resuspended in cold PBS or Versene. Then, single cell suspensions were labelled with a viability marker (Zombie Violet; BioLegend, 423114) at 4°C for 15 min. PBS alone or PBS/2mM EDTA (Supelco, 1.08418)/0.5% BSA was used as buffer solution.

For extracellular staining, cells were directly incubated with antibodies or the corresponding isotypes, diluted at the appropriate concentration in buffer solution, at 4°C for 30 min. Antibodies are listed in Table S2.

For intracellular staining, cells were fixed, permeabilized and stained with the appropriate antibodies (direct or indirect staining) and the Transcription Factor Staining Buffer Set (Miltenyi Biotec, 130-122-981) following the manufacturer’s instructions or with 4% paraformaldehyde solution and BD Phosflow Perm/Wash buffer (BD Biosciences, 557885), as previously described (26). A FACSCanto flow cytometry system and FloJow (BD Biosciences) were used for data acquisition and analysis.

### Reverse Transcription-Quantitative Polymerase Chain Reaction (RT-qPCR)

RNA was extracted from cells using the QIAshredder Kit (QIAGEN, Redwood city, CA, USA) or the RNeasy Mini Kit (QIAGEN, Redwood city, CA, USA) according to the manufacturer’s instructions. RNA was quantified with a NanoDrop spectrophotometer (Thermo Fisher Scientific, Waltham, United States). RT-qPCR was performed using the Luna Universal One-Step RT-qPCR Kit (New England Biolabs, Ipswich, MA, USA) and a LightCycler 480 system (Roche Diagnostics). Primers are listed in Table S3. Each sample was amplified in triplicate and gene expression levels were normalized to the expression of the housekeeping gene *GAPDH* (ΔCt). Relative gene expression was expressed as fold-change using the formula: Log10(2^-ΔCt^+1e-6). The lowest value of the negative control was then subtracted.

### Cilia beating imaging

For indirect assessment of cilia beating, fluorescent microbead suspensions were applied to the apical surface of iALI cultures, and their movement was recorded using a THUNDER Imaging system mounted on a stereomicroscope (Leica Microsystems). Microbead trajectories were reconstructed in Fiji (ImageJ) and then analysed and visualized using R scripts (ggplot2 library).

### Replicates

The cell graft experiment was performed 28 times by multiple operators over 3.5 years, representing a total number of 528 iALI wells. Among them, 436 iALI wells were derived from the parental Hy03 iPSC line and 92 iALI wells were derived from the Hy03 MCIDAS^-/-^ iPSC line.

### Statistics

Differences between independent groups were assessed using the Mann-Whitney U test. This non-parametric test was chosen because the data did not meet the assumptions of normal distribution (Shapiro-Wilk test). A p-value < 0.05 was considered statistically significant.

## Results

### vAFE cells derived from iPSCs can differentiate into a functional airway epithelium

The parental Hy03 and Hy03 GFP iPSC lines successfully differentiated into functional airway epithelium in iALI, as previously described by our team (Fig.1A) (26). iPSCs differentiated into lung progenitors (vAFE cells) in 7 days. They passed through the different embryonic stages of lung development (anterior primitive streak at day 2, definitive endoderm at day 3, anterior foregut endoderm and vAFE at days 4-7). Of note, this differentiation protocol does not require any cell sorting step. At day 7, lung progenitors were characterized by strong expression of the NKX2.1 transcription factor, assessed by RT-qPCR (∼5-10-fold more than iPSCs) and flow cytometry (∼80% of cells in culture) (Fig. 1B,C). At the vAFE stage, RT-qPCR confirmed the very low number of pluripotent stem cells (*SOX2*+) and of cells belonging to the neuronal (PAX6), intestinal (HNF4A) and thyroid (PAX8) lineages (Fig. S1).

**Figure 1.**
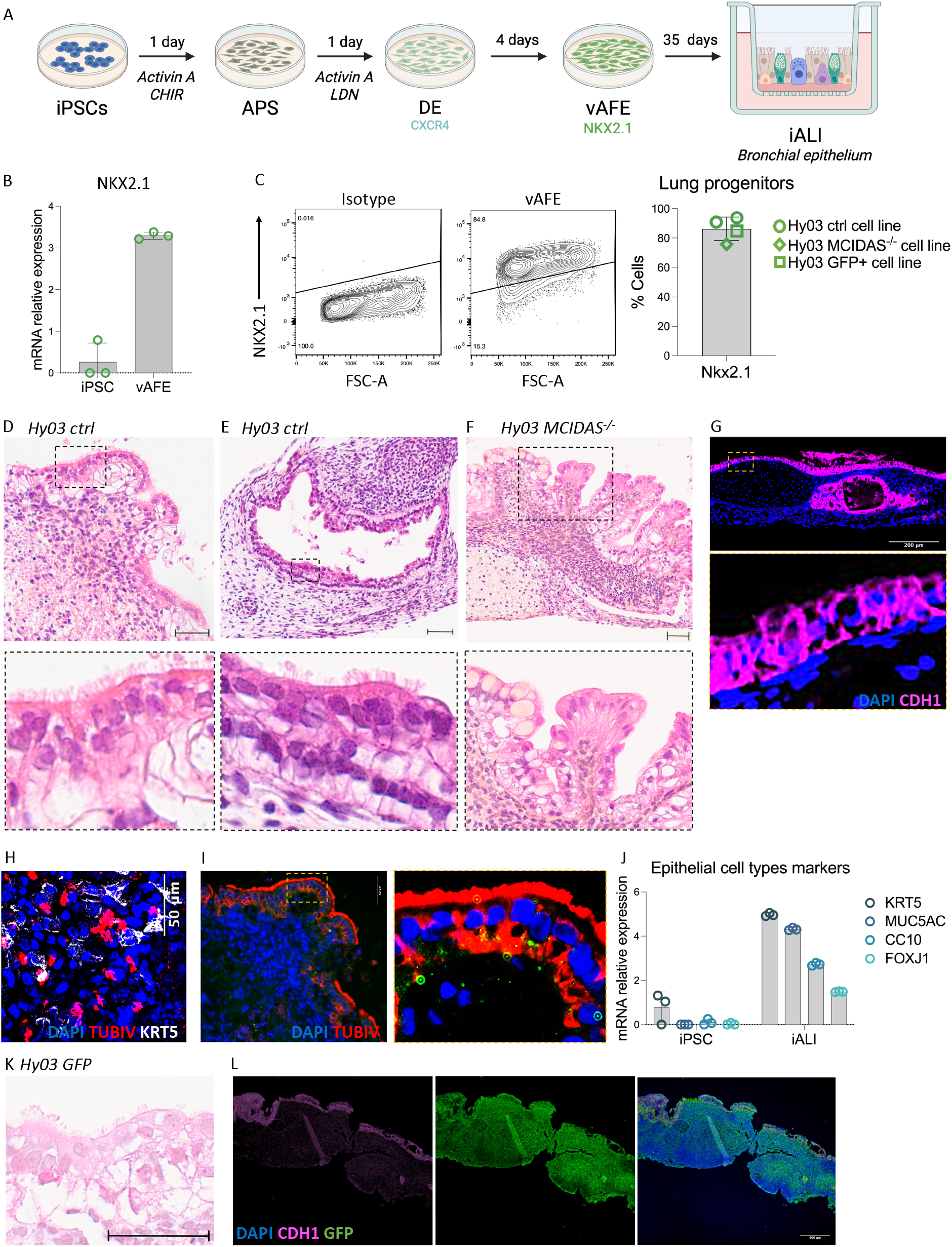
Lung progenitors (vAFE) derived from iPSCs can differentiate into a mature bronchial epithelium (iALI). (A) Schematic summary of the protocol of directed differentiation of iPSCs into an iALI bronchial epithelium. (B) mRNA expression level of bronchial progenitor (vAFE stage) markers assessed by RT-qPCR (experiment reproduced in triplicate). The data shown are for progenitors differentiated from the Hy03 GFP iPSC line. (C) Flow cytometry quantification of NKX2.1 expression in bronchial progenitors (vAFE stage) (n=4 independent experiments). (D, E, F) Representative images of iALI sagittal sections stained with haematoxylin-eosin-saffron after differentiation from the Hy03 and Hy03 MCIDAS^-/-^ iPSC lines (day 35) (scale-bar = 50μm). Note the absence of cilia in the Hy03 MCIDAS^-/-^ iALI culture. (G) Representative image of E-cadherin (CDH1; pink) expression in an iALI sagittal section after differentiation from the Hy03 iPSC line (day 35). (H) Representative image of tubulin-IV (red) and KRT5 (white) expression at the apical surface of an iALI culture after differentiation from the Hy03 iPSC line (day 35). (I) Representative image of tubulin-IV (red) expression in an iALI sagittal section after differentiation from the Hy03 iPSC line (day 35). (J) mRNA expression analysis by RT-qPCR of the indicated epithelial cell markers in iALI after differentiation from the Hy03 iPSC line (day 35) (experiment reproduced in triplicate). (K) Representative image of haematoxylin-eosin-saffron staining in iALI sagittal sections after differentiation from the Hy03 GFP IPSC line (day 35) (scale bar = 50μm). (L) Representative images of GFP (green) and E-cadherin (pink) expression in an iALI sagittal section after differentiation from the Hy03 GFP iPSC line (day 35). APS: Anterior Primitive Streak; DE: Definitive Endoderm; vAFE: ventralized Anterior Foregut Endoderm.

Then, lung progenitors (day 7) were seeded on Transwell™ plates and expanded for approximately 35 days in differentiation medium (Fig.1A). At day 42, a structured and polarized ciliated epithelium could be observed (Fig.1D). Occasionally, rounded bronchus-like epithelial structures were found buried under the surface (Fig.1E). Immunofluorescence analysis was used to confirm the epithelial organization and identity. This demonstrated that E-cadherin (CDH1)-positive epithelial cells lined the surface and were supported by underlying E-cadherin–negative mesenchymal cells (Fig.1G). The main different cell types of the adult respiratory epithelium were identified by immunofluorescence staining and RT-qPCR (Fig.1H-J): KRT5-expressing basal cells, MUC5AC- and SCGB1A1/CC10-expressing secretory cells (goblet and club cells) and tubulin IV (TubIV)- and FOXJ1-expressing ciliated cells. Cilia beating was analysed by microscopic video (Video S1). Similarly, the Hy03 GFP iPSCs differentiated readily into an E-cadherin+ multiciliated epithelium that lined the surface of E-cadherin-negative mesenchymal cells (Fig.1K,L).

The Hy03 MCIDAS^-/-^ iPSC line also could be differentiated into airway epithelium in iALI (Fig. 1C). However, atrophic cilia were observed in these cultures (Fig.1F compared with Fig. 1D and E for Hy03 iPSCs).

### Early lung progenitor cells can integrate into mature iALI lung epithelium

Effective cell grafting may require “airway conditioning,” a process that involves the deliberate disruption and removal of the existing bronchial epithelium to allow the durable engraftment of gene-corrected cells. As a proof of concept, first trypsin-based conditioning of the parental Hy03 iPSC-derived iALI before bronchial progenitor graft was tested (Fig. 2A). The conditioning efficiency was assessed by measuring the de-epithelialized zone length on sagittal iALI sections after different times of incubation with trypsin. Trypsin (0.25%) induced a shedding of 10% and 50% of the epithelium after incubation for 5 and 25 minutes, respectively (Fig. 2B).

**Figure 2.**
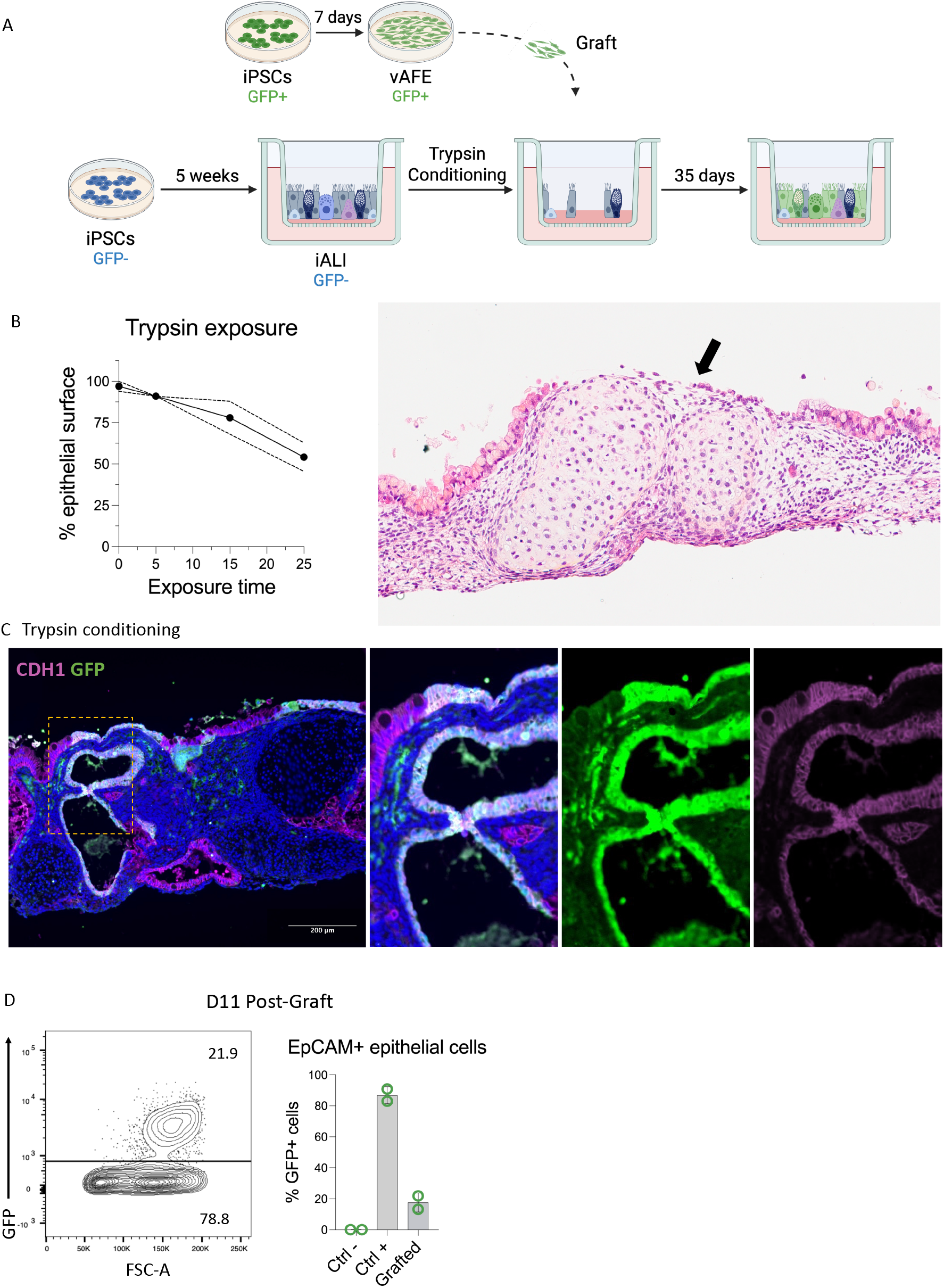
vAFE cells can engraft in a bronchial epithelial model after trypsin conditioning. (A) Schematic summary of the engraftment experiment on iALI. (B) Epithelial shedding in function of trypsin exposure time (left). The length of the de-epithelized surface (arrow in right panel) relative to the total surface of the haematoxylin-eosin-saffron stained sagittal section was measured using the NDP view software (n=2 independent experiments). (C) Immunofluorescence analysis of the integration of engrafted GFP+ progenitors (green) in the E-cadherin+ epithelial structure (CDH1; pink) in a Hy03 iALI engrafted with Hy03 GFP vAFE cells at day 35 post-graft. The recipient iALI was conditioned with 0.25% trypsin for 15 minutes. (D) Flow cytometry quantification of EpCAM+ GFP+ cells at day 11 post-graft to quantify the epithelial integration and differentiation of GFP+ progenitors (n = 2 independent experiments). Ctrl-: parental Hy03 iPSC-derived iALI; Ctrl+: Hy03 GFP iPSC-derived iALI; GFP+: Hy03 GFP iPSC line; GFP-: parental Hy03 iPSC line.

Then, lung progenitors (vAFE stage) derived from the Hy03 GFP iPSC line (GFP+ progenitors hereafter) were seeded on the conditioned apical surface of mature iALI cultures from the Hy03 iPSC line at week 5 of differentiation. Grafted iALI cultures were maintained in culture for another 35 days. At day 70, GFP+ cells largely contributed to the epithelium of the grafted iALI, as indicated by E-cadherin and GFP co-staining (Fig. 2C). The integration and differentiation of GFP+ progenitors was assessed by flow cytometry analysis of EpCAM+ GFP+ cells at day 11 post-graft (Fig. 2D): ∼20% of the sorted EpCAM+ cells were GFP+ (n = 2 independent experiments). Collectively, these assays suggest that vAFE cells are good candidates for epithelial cell replacement because they successfully integrated and differentiated into airway epithelium.

### Conditioning of iALI before cell grafting

Airway conditioning remains an unmet requirement for the successful implementation of cell replacement. Therefore, besides trypsin, other enzymatic and chemical strategies for conditioning were investigated (Fig. 3A)(Table S1). The balance between efficacy and toxicity was assessed by quantifying cell detachment, cell viability and engraftment yield. Comparison of the number of detached cells after exposure to the different compounds showed that all tested strategies, except polidocanol, induced epithelial shedding compared with control (Pneumacult™ ALI Basal Medium) (Fig. 3B; n=2 to 8 independent experiments). Although enzymatic solutions were more effective in detaching cells, chemical conditioning approaches (EDTA, CHAPS, polidocanol) were further tested because of their potentially lower production costs and likely reduced risk of allergic side effects. Cell viability was assessed by flow cytometry after Zombie Violet dye staining at day 7 after chemical conditioning (Fig. 3C). Polidocanol was associated with significant cell mortality even at a relatively low concentration (0.5%). The toxicity of polidocanol compared with the other regimens was confirmed at day 35 post-graft (Fig. 3D). As EDTA offers multiple advantages, including low cost, ease of use, acceptable cell detachment efficiency, and good cell viability at day 7 and day 35, this approach was selected for further investigation. At day 35 after grafting of GFP+ progenitors in EDTA pre-conditioned Hy03-derived iALI, immunofluorescence staining confirmed the integration of GFP+ cells in the epithelium (E-cadherin-positive cells) (Fig. 3E), similarly to what previously observed with trypsin.

**Figure 3.**
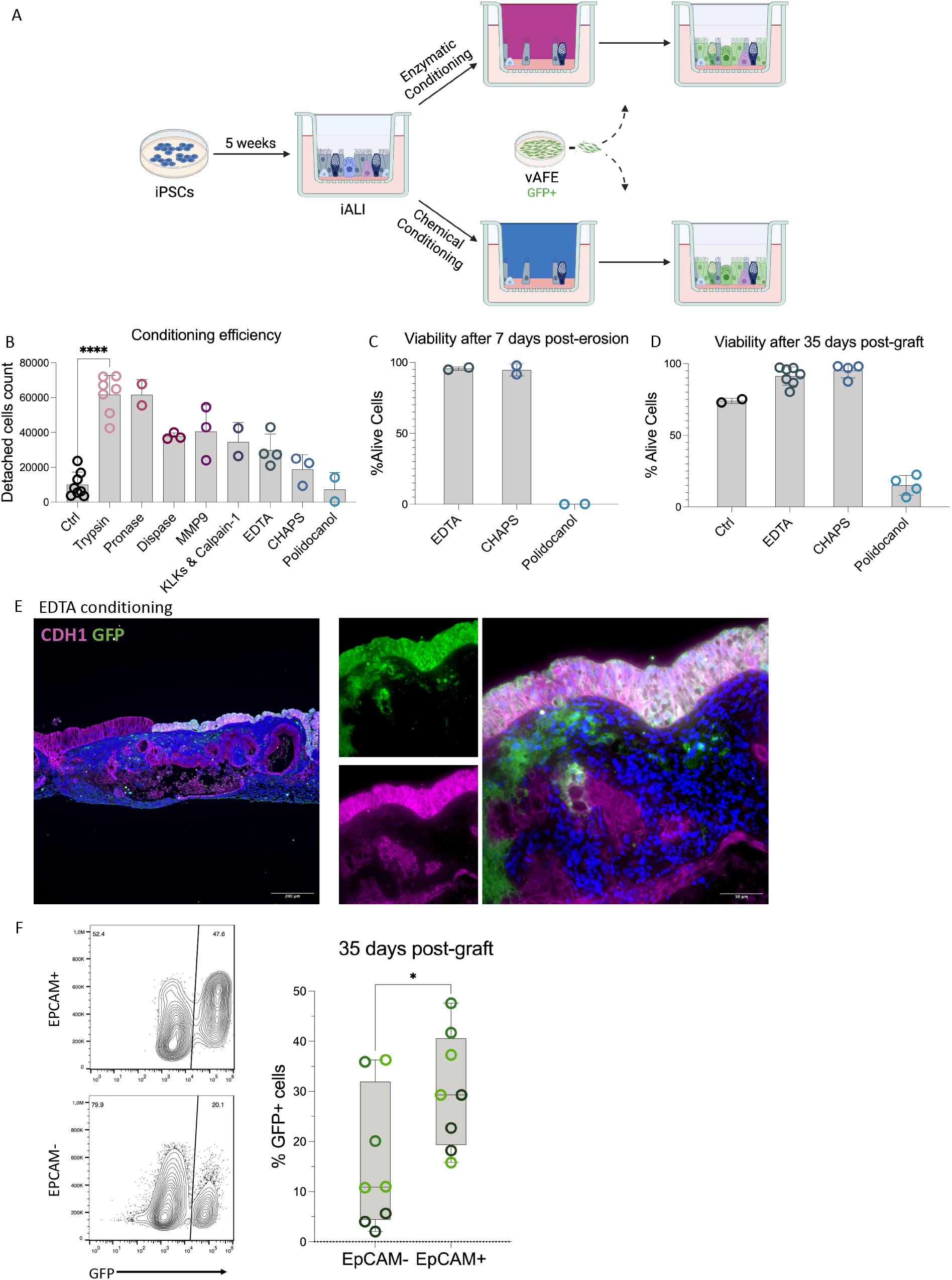
Balance between efficiency and toxicity of enzymatic and chemical compounds used for the preconditioning step to promote cell engraftment. (A) Schematic summary of the engraftment experiment on iALI after different conditioning strategies. (B) Number of detached cells after exposure to the different compounds (n = 2 to 8 independent experiments). (C,D) Flow cytometry quantification of living cells in iALI at day 7 (C) and 35 (D) after exposure to the indicated chemical compounds (n = 2 to 7 independent experiments). (E) Immunofluorescence analysis of the integration of engrafted GFP+ progenitors (green) in the E-cadherin+ epithelial structure (CDH1; pink) of the iALI after EDTA conditioning at day 35 post-graft. (F) Quantification of GFP+ cells in the mesenchymal (EpCAM-) and in the epithelial (EpCAM+) cell populations in iALI at day 35 post-graft after EDTA conditioning (each point is representative for an iALI well, and each colour of a different engraftment experiment).

Engrafted GFP+ cells were quantified by flow cytometry at day 35 post-graft. Co-staining of EpCAM allowed the quantification of GFP+ cells integrated in the epithelial and non-epithelial compartments (3 independent assays with 2 to 3 replicates each). The success of epithelial cell replacement (i.e. percentage of GFP+ cells among all EpCAM+ cells) ranged from 15 to 48% (Fig. 3F). Moreover, up to 37% of non-epithelial EpCAM-cells expressed GFP, suggesting that vAFE cells can integrate also into the mesenchymal layer, albeit with lower efficiency than in the epithelial layer. In addition, the engraftment of GFP+ progenitors on non-eroded iALI also was successful, reaching similar levels of epithelial cell integration by flow cytometry and immunofluorescence analysis (Fig. S2).

### Engrafted progenitors mature into adult epithelial cell types and ensure functional recovery of PCD iALI

At day 35 post-graft, the fate of engrafted GFP+ progenitors was assessed to evaluate their long-term persistence and potential contribution to functional repair (Fig. 4A). GFP staining of paraffin sections from recipient Hy03-derived iALI cultures allowed identifying the GFP+ ciliated epithelium (brown staining, Fig. 4B). Moreover, immunohistochemical analysis showed that GFP+ cells had differentiated into ciliated cells (tubulin IV+) and goblet cells (MUC5AC+) (Fig. 4C,D). Flow cytometry analysis of cells co-stained with anti-GFP and -KRT5 antibodies showed that almost 10% of the engrafted GFP+ cells were KRT5+ basal cells (Fig. 4E).

**Figure 4.**
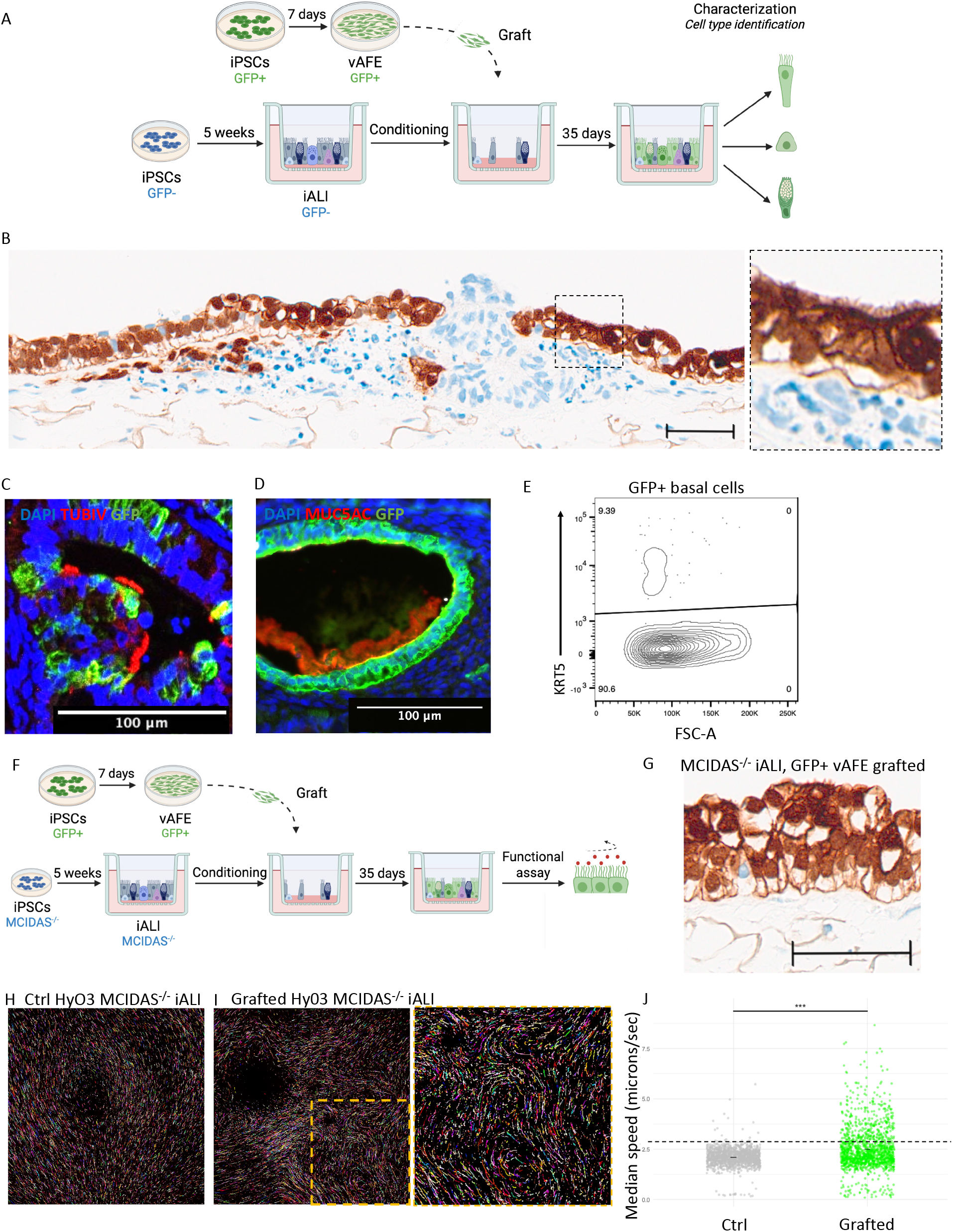
Maturation of the engrafted lung progenitors and functional repair. (A) Schematic summary of the cell fate characterization after engraftment. (B) Representative image of a sagittal section from a parental Hy03-derived iALI at day 35 after graft of Hy03 GFP iPSC-derived progenitor cells after EDTA conditioning. Immunohistochemical staining was performed using an anti-GFP antibody. (C, D) Immunofluorescence characterization of cells derived from engrafted GFP+ lung progenitors (green) that have differentiated into tubulin-IV+ ciliated cells (red, C) and MUC5AC+ goblet cells (red, D) (day 35 post-graft). (E) Flow cytometry quantification of basal cells derived from engrafted GFP+ lung progenitors identified by co-staining with anti-KRT5 and -GFP antibodies at day 35 post-graft (n=1). (F) Schematic summary of the experiment to assess whether healthy GFP+ progenitor engrafting can lead to cilia beating recovery in iALI derived from the Hy03 MCIDAS^-/-^ iPSC line (PCD model). (G) Representative sagittal section of an Hy03 MCIDAS^−^/^−^ derived epithelium at day 35 post-grafting, showing GFP^+^ cells detected by immunohistochemical staining with an anti-GFP antibody. The recipient iALI was conditioned with EDTA (amount, time) and Hy03 GFP vAFE cells were used as donor cells. (H and I) Assessment of cilia function by monitoring the movement of fluorescent beads on the epithelium surface. (J) Quantification of the bead speed on a non-grafted Hy03 MCIDAS^-/-^ iALI (Ctrl) and in a MCIDAS^-/-^ iALI grafted with Hy03 GFP iPSC-derived vAFE cells (day 35 post-graft) (n=1 to 2).

As an *in vitro* proof of concept of the potential of this strategy in PCD, GFP+ progenitors were grafted in iALIs derived from the Hy03 MCIDAS^-/-^ iPSC line at week 5 of differentiation and after conditioning with EDTA (Fig. 4F). At day 35 post-graft, ciliated cells derived from engrafted GFP+ cells were detected by immunohistochemistry (Fig.4G). Cilia movement tracking using fluorescent microbeads (Video S2) showed the absence of any significant bead movement in control (non-grafted) Hy03 MCIDAS^-/-^ iALI (Fig. 4H). Conversely, bead movement was recorded in Hy03 MCIDAS^-/-^ iALI grafted with GFP+ progenitors (Video S3). In some areas, these beads followed apparently organized trajectories, suggesting the recovery of a coordinated and functional ciliary beating activity (Fig. 4I). Moreover, in the grafted condition, the speed of the beads was significantly increased compared with the non-grafted condition (Fig. 4J). These findings indicate that engrafting healthy GFP+ vAFE cells to replace Hy03 MCIDAS^-/-^ derived cells leads to successful homing and replacement of MCIDAS^-/-^ cells by normally beating GFP+ ciliated cells that can generate mucociliary motion.

## Discussion

This study investigated the potential of vAFE cells as early lung progenitors to replace and repair a mature bronchial epithelium in view of developing a combined gene and cell therapy strategy for treating monogenic lung diseases. Early lung progenitors can be derived from iPSCs following a differentiation protocol that mimic embryonic development. We accumulated compelling evidence that these progenitors can differentiate into the most frequent cell types of the adult bronchial epithelium, such as basal and ciliated cells. This was achieved not at the cost of higher rates of contamination by other tissue cell types. These characteristics make them credible candidates for future therapeutic applications. Then, we assessed their ability to engraft a preconditioned iALI lung epithelium. Importantly, no cell sorting was necessary before transplantation. The lung progenitors integrated the model, especially in the epithelial compartment at rates up to 48% at day 35 post-graft. However, these early progenitors may have high and hard-to-control proliferation rates. Moreover, they might have a less well-established airway signature associated with higher plasticity, increasing the risk of ‘contamination’/differentiation toward undesirable tissues (e.g. liver, thyroid). More studies are needed to explore these issues.

To optimize progenitor engraftment and repair, we compared different conditioning regimens. Enzymatic conditioning yielded promising results; however, its clinical application presents several challenges (toxicity of non-specific enzymes, cost of producing specific enzymes, immune response). Conversely, chemical conditioning offers distinct advantages in terms of production, although the safety of the considered solutions must be assessed *in vivo*. Indeed, polidocanol showed high cell toxicity, whereas EDTA and CHAPS represent promising alternatives. As EDTA safety has already been demonstrated in humans and is relatively easy to use (34,35), we focused on this strategy for iALI conditioning before lung progenitor transplantation. We showed that early lung progenitors integrated the iALI after EDTA-induced epithelial erosion. Surprisingly, we observed similar engraftment rates with and without conditioning in our cell model (Fig. S2). However, *in vivo*, the mucosal barrier (despite ineffective mucociliary clearance in patients with PCD) and efficient cough, possibly in an infectious environment, would limit cell engraftment, highlighting the potential need for conditioning. Indeed, conditioning may facilitate the access of transplanted cells to stem cell niches, including the ones harbouring the highly resistant Hillock stem cells that are essential for long-term self-renewal of the corrected epithelium and sustained therapeutic efficacy.

Restoration of the microbead movement at the apical surface of the grafted PCD iALI demonstrates the recovery of the mucociliary clearance function after vAFE cell engraftment. Functional restauration of ciliary beating in physiological conditions requires that at least 15% to 30% of epithelial cells express functional cilia (36). In the present study, we observed engraftment rates ranging from 15% to 45% in the epithelium compartment. If comparable efficiencies could be achieved after *in vivo* translation, and if ∼50% of engrafted cells would differentiate into ciliated cells, a minimum of 30% of the epithelial compartment would need to be derived from donor cells to achieve functional repair. Our results show that this is achievable *in vitro*, even before optimization of the engraftment process, which will be pursued in future studies.

### Limitations

We acknowledge several limitations. Engraftment analyses revealed that a subset of transplanted cells localized beneath the epithelial compartment. The identity and fate of these cells remain to be fully elucidated. One hypothesis is that they originate from the small NKX2.1-negative subpopulation present at the vAFE stage, raising the possibility of heterogeneity in the engraftment competence or lineage potential. Future studies that combine lineage tracing and refined phenotypic profiling will be necessary to determine whether these cells represent transient intermediates, mislocalized progenitors, or a distinct population. Although cell sorting was not required to achieve functional rescue in the current experimental context, selective enrichment strategies may become relevant as translational objectives evolve.

The observation of epithelial mosaicism, with juxtaposed GFP-positive and GFP-negative regions, is consistent with the clonal expansion of engrafted cells and suggests a partial replacement of endogenous basal stem cells. This is supported by the emergence of differentiated epithelial lineages, including ciliated and mucus-secreting cells, that correspond to the expected airway epithelial hierarchies. The absence of club cells in our cultures could be attributed to pharmacological inhibition of the Notch pathway by DAPT, underscoring the importance of culture conditions in shaping lineage outcomes.

An important and somewhat unexpected finding was that engraftment could occur in the absence of deliberate epithelial injury. This observation challenges the prevailing assumptions that conditioning-induced damage is a prerequisite for donor cell integration. The mechanisms underlying injury-independent engraftment remain unclear, but may involve physiological processes, such as cell extrusion in overcrowded epithelial regions or competitive niche replacement. Dissecting these mechanisms will be essential for the rational optimization of engraftment strategies and may inform approaches that minimize epithelial injury in future therapeutic applications.

Despite these encouraging findings, significant challenges remain before application in patients can be envisioned. Clinical translation will require validation *in vivo*, ideally in large animal models that more closely recapitulate human airway anatomy and physiology, as well as compliance with the Good Manufacturing Practice (GMP) standards for cell production. Additionally, long-term durability of the grafted cells, epithelial cell turnover dynamics, and immune response issues will need to be rigorously assessed.

## Conclusions

In summary, this study establishes that vAFE cells can engraft into human airway epithelial models and restore functional ciliary activity in a disease-relevant context. Conditioning strategies may enhance engraftment efficiency; however, their necessity in physiological *in vivo* conditions remains to be determined. Together, these findings provide a foundation for preclinical investigations and represent an important step toward cell-based regenerative therapies for genetic airway diseases.

## List of abbreviations

APS: Anterior Primitive Streak
CDH1: E-cadherin
DE: Definitive Endoderm
GFP: Green Fluorescent Protein
iALI: iPSC-derived bronchial epithelium in Air-Liquid Interface
iPSC: induced Pluripotent Stem Cell
KRT5: Cytokeratine 5
PCD: Primary Ciliary Dyskinesia
RT: Room Temperature
RT-qPCR: Reverse Transcription-Quantitative Polymerase Chain Reaction
TubIV: Tubulin IV
vAFE: ventral Anterior Forgut Endoderm

## Ethics approval

The UHOMi002-A human iPSC line was derived from a healthy, non-smoking donor. The study received approval from the regional ethics committee (CPP Sud Med IV; ID-RCB: 2017-A00252-51). The University Hospital of Montpellier (France) acted as the study sponsor, and written informed consent was obtained from the donor.

## Availability of data and materials

The datasets used and/or analysed during the current study are included in this published article (and its supplementary files) or available from the corresponding author on reasonable request.

## Authors’ contributions

Conceptualization and supervision: C.B., A.B., S.A. and J.D.V.; resources: A.B. and J.D.V.; data analysis and interpretation: C.B., A.B., S.A. and J.D.V; data collection and/or assembly: C.B., A.C., F.F., M.N., C.U., A.N., J.M., L.M., F.R., L.Y., A.P., Q.B., I.V., S.A., A.B. and J.D.V.; writing-original draft: C.B., A.B., and J.D.V.; writing-review and editing: C.B., A.B., S.A. and J.D.V.; scientific and technical support: all authors listed.

## Acknowledgements

We thank the Biocampus Montpellier imaging facility MRI, member of the national infrastructure France-BioImaging (https://ror.org/01y7vt929) supported by the French National Research Agency (ANR-24-INBS-0005 FBI BIOGEN), and the “Réseau d’Histologie Expérimentale de Montpellier” (RHEM) facility, which is supported by SIRIC Montpellier Cancer Grant INCa_Inserm_DGOS_12553, the European regional development foundation and the Occitanie region (FEDER-FSE 2014-2020 Languedoc Roussillon), for histology sample preparation. We also acknowledge the support of Immun4Cure IHU “Institute for innovative immunotherapies in autoimmune diseases” (France 2030 / ANR-23-IHUA-0009). We thank Yoan Arribat’s group for providing access to their stereomicroscope.

## Figure legends

Figure S1. mRNA expression level analysis by RT-qPCR of the indicated markers of contaminant lineages (PAX6, PAX8, HNF4A) and pluripotent stem cells (SOX2) in the Hy03 GFP iPSC line and the derived vAFE (experiment reproduced in triplicate).

Figure S2. (A) Immunofluorescence analysis of the integration of engrafted GFP+ progenitors (green) in the E-cadherin positive epithelial structure (CDH1; pink) of an Hy03-derived iALI without conditioning. Scale bar = (B) Comparison of GFP+ cell engraftment in the epithelial (EpCAM+) compartment of Hy03-derived iALIs with and without (Ctrl) EDTA conditioning (n=8 to 9 independent experiments).

Table S1. Compounds used for the conditioning step before cell grafts. In the table change commas to points (ex. 0,25% to 0.25%). Same comment also for the Tm in Table S3.

Table S2. Antibodies used for flow cytometry (FC) and immunofluorescence (IF) experiments.

Table S3. Primers used for RT-qPCR.

Video S1. Video showing cilia beating in the bronchial epithelium.

Video S2. Video showing the movement of fluorescent microbeads on the bronchial epithelium surface of an Hy03 MCIDAS^−^/^−^ derived epithelium.

Video S3. Video showing the movement of fluorescent microbeads on the bronchial epithelium surface of an Hy03 MCIDAS^−^/^−^ derived epithelium at day 35 post-grafting.

## References

1. Horani A, Ferkol TW. Understanding Primary Ciliary Dyskinesia and Other Ciliopathies. J Pediatr. 2021 Mar;230:15–22.e1.

2. Horani A, Ferkol TW. Advances in the Genetics of Primary Ciliary Dyskinesia. Chest. 2018 Sep;154(3):645–52.

3. Mianné J, Ahmed E, Bourguignon C, Fieldes M, Vachier I, Bourdin A, et al. Induced Pluripotent Stem Cells for Primary Ciliary Dyskinesia Modeling and Personalized Medicine. Am J Respir Cell Mol Biol. 2018 Dec;59(6):672–83.

4. Keicho N, Morimoto K, Hijikata M. The challenge of diagnosing primary ciliary dyskinesia: a commentary on various causative genes and their pathogenic variants. J Hum Genet. 2023 Aug;68(8):571–5.

5. Wee WB, Kinghorn B, Davis SD, Ferkol TW, Shapiro AJ. Primary Ciliary Dyskinesia. Pediatrics. 2024 Jun 1;153(6):e2023063064.

6. Bukowy-Bieryllo Z, Witt M, Zietkiewicz E. Perspectives for Primary Ciliary Dyskinesia. Int J Mol Sci. 2022 Apr 8;23(8):4122.

7. Boon M, Wallmeier J, Ma L, Loges NT, Jaspers M, Olbrich H, et al. MCIDAS mutations result in a mucociliary clearance disorder with reduced generation of multiple motile cilia. Nat Commun. 2014 Jul 22;5:4418.

8. Lai M, Pifferi M, Bush A, Piras M, Michelucci A, Di Cicco M, et al. Gene editing of DNAH11 restores normal cilia motility in primary ciliary dyskinesia. J Med Genet. 2016 Apr;53(4):242–9.

9. Ostrowski LE, Yin W, Patel M, Sechelski J, Rogers T, Burns K, et al. Restoring ciliary function to differentiated primary ciliary dyskinesia cells with a lentiviral vector. Gene Ther. 2014 Mar;21(3):253–61.

10. Driskell RA, Engelhardt JF. Current status of gene therapy for inherited lung diseases. Annu Rev Physiol. 2003;65:585–612.

11. Paff T, Omran H, Nielsen KG, Haarman EG. Current and Future Treatments in Primary Ciliary Dyskinesia. Int J Mol Sci. 2021 Sep 11;22(18):9834.

12. Allan KM, Farrow N, Donnelley M, Jaffe A, Waters SA. Treatment of Cystic Fibrosis: From Gene-to Cell-Based Therapies. Front Pharmacol. 2021 Mar 16;12:639475.

13. Berical A, Lee RE, Randell SH, Hawkins F. Challenges Facing Airway Epithelial Cell-Based Therapy for Cystic Fibrosis. Front Pharmacol. 2019 Feb 8;10:74.

14. Adams TS, Marlier A, Kaminski N. Lung Cell Atlases in Health and Disease. Annu Rev Physiol. 2023 Feb 10;85(1):47–69.

15. He P, Lim K, Sun D, Pett JP, Jeng Q, Polanski K, et al. A human fetal lung cell atlas uncovers proximal-distal gradients of differentiation and key regulators of epithelial fates. Cell. 2022 Dec;185(25):4841–4860.e25.

16. Rock JR, Onaitis MW, Rawlins EL, Lu Y, Clark CP, Xue Y, et al. Basal cells as stem cells of the mouse trachea and human airway epithelium. Proc Natl Acad Sci USA. 2009 Aug 4;106(31):12771–5.

17. Zuo W, Zhang T, Wu DZ, Guan SP, Liew AA, Yamamoto Y, et al. p63+Krt5+ distal airway stem cells are essential for lung regeneration. Nature. 2015 Jan 29;517(7536):616–20.

18. Ghosh M, Ahmad S, White CW, Reynolds SD. Transplantation of Airway Epithelial Stem/Progenitor Cells: A Future for Cell-Based Therapy. Am J Respir Cell Mol Biol. 2017 Jan;56(1):1–10.

19. Lin B, Shah VS, Chernoff C, Sun J, Shipkovenska GG, Vinarsky V, et al. Airway hillocks are injury-resistant reservoirs of unique plastic stem cells. Nature. 2024 May;629(8013):869–77.

20. Mou H, Vinarsky V, Tata PR, Brazauskas K, Choi SH, Crooke AK, et al. Dual SMAD Signaling Inhibition Enables Long-Term Expansion of Diverse Epithelial Basal Cells. Cell Stem Cell. 2016 Aug 4;19(2):217–31.

21. Cerneckis J, Cai H, Shi Y. Induced pluripotent stem cells (iPSCs): molecular mechanisms of induction and applications. Sig Transduct Target Ther. 2024 Apr 26;9(1):112.

22. Takahashi K, Yamanaka S. Induction of pluripotent stem cells from mouse embryonic and adult fibroblast cultures by defined factors. Cell. 2006 Aug 25;126(4):663–76.

23. Yamanaka S. Pluripotent Stem Cell-Based Cell Therapy-Promise and Challenges. Cell Stem Cell. 2020 Oct 1;27(4):523–31.

24. Goecke T, Ius F, Ruhparwar A, Martin U. Unlocking the Future: Pluripotent Stem Cell-Based Lung Repair. Cells. 2024 Apr 5;13(7):635.

25. Mianné J, Nasri A, Van CN, Bourguignon C, Fieldès M, Ahmed E, et al. CRISPR/Cas9-mediated gene knockout and interallelic gene conversion in human induced pluripotent stem cells using non-integrative bacteriophage-chimeric retrovirus-like particles. BMC Biol. 2022 Dec;20(1):8.

26. Ahmed E, Fieldes M, Bourguignon C, Mianné J, Petit A, Vernisse C, et al. Differentiation of human induced pluripotent stem cells into functional airway epithelium [Internet]. Bioengineering; 2020 Nov [cited 2024 Mar 6]. Available from: http://biorxiv.org/lookup/doi/10.1101/2020.11.29.400358

27. Calvert BA, Ryan AL. Application of iPSC to Modelling of Respiratory Diseases. In: Turksen K, editor. Cell Biology and Translational Medicine, Volume 7 [Internet]. Cham: Springer International Publishing; 2019 [cited 2024 Dec 26]. p. 1–16. (Advances in Experimental Medicine and Biology; vol. 1237). Available from: https://link.springer.com/10.1007/5584_2019_430

28. Humbert MV, Spalluto CM, Bell J, Blume C, Conforti F, Davies ER, et al. Towards an artificial human lung: modelling organ-like complexity to aid mechanistic understanding. Eur Respir J. 2022 Dec;60(6):2200455.

29. McCauley KB, Hawkins F, Serra M, Thomas DC, Jacob A, Kotton DN. Efficient Derivation of Functional Human Airway Epithelium from Pluripotent Stem Cells via Temporal Regulation of Wnt Signaling. Cell Stem Cell. 2017 Jun 1;20(6):844–857.e6.

30. Vazquez-Armendariz AI, Tata PR. Recent advances in lung organoid development and applications in disease modeling. Journal of Clinical Investigation. 2023 Nov 15;133(22):e170500.

31. Hawkins F, Kramer P, Jacob A, Driver I, Thomas DC, McCauley KB, et al. Prospective isolation of NKX2-1–expressing human lung progenitors derived from pluripotent stem cells. Journal of Clinical Investigation. 2017 May 2;127(6):2277–94.

32. Longmire TA, Ikonomou L, Hawkins F, Christodoulou C, Cao Y, Jean JC, et al. Efficient Derivation of Purified Lung and Thyroid Progenitors from Embryonic Stem Cells. Cell Stem Cell. 2012 Apr;10(4):398–411.

33. Fieldes M, Ahmed E, Bourguignon C, Mianné J, Martin M, Arnould C, et al. Generation of the induced pluripotent stem cell line UHOMi002-A from peripheral blood mononuclear cells of a healthy male donor. Stem Cell Research. 2020 Dec;49:102037.

34. Lanigan RS, Yamarik TA. Final Report on the Safety Assessment of EDTA, Calcium Disodium EDTA, Diammonium EDTA, Dipotassium EDTA, Disodium EDTA, TEA-EDTA, Tetrasodium EDTA, Tripotassium EDTA, Trisodium EDTA, HEDTA, and Trisodium HEDTA. Int J Toxicol. 2002 Oct;21(2_suppl):95–142.

35. Wang G, Zabner J, Deering C, Launspach J, Shao J, Bodner M, et al. Increasing Epithelial Junction Permeability Enhances Gene Transfer to Airway Epithelia In Vivo. Am J Respir Cell Mol Biol. 2000 Feb 1;22(2):129–38.

36. Loges NT, Marthin JK, Raidt J, Olbrich H, Höben IM, Cindric S, et al. A range of 30–62% of functioning multiciliated airway cells is sufficient to maintain ciliary airway clearance. Eur Respir J. 2024 Oct;64(4):2301441.

