## Supplemental figures, tables and videos for "Ventral anterior foregut endoderm cells as progenitors for airway epithelial cell replacement in primary ciliary dyskinesia"

S1

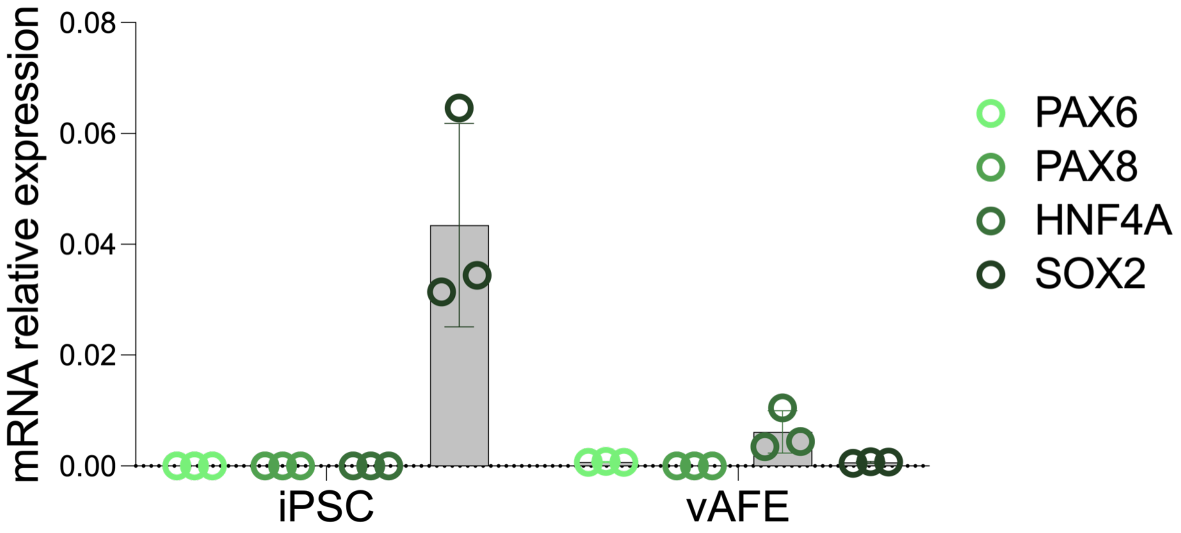

mRNA relative expression
0.08
0.06
0.04
0.02
0.00
iPSC
vAFE
PAX6
PAX8
HNF4A
SOX2


S2

A

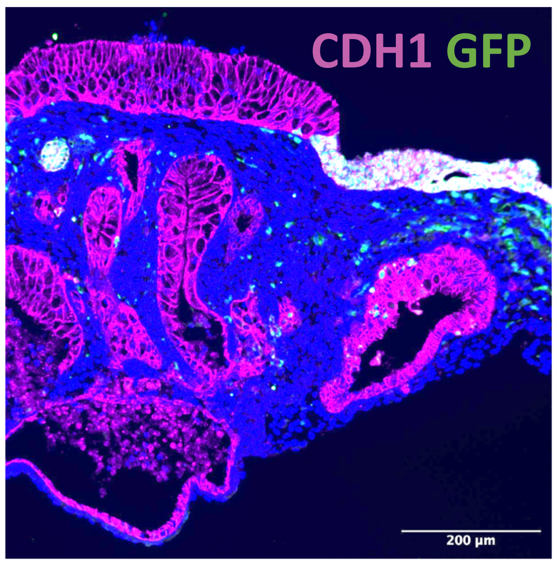

CDH1 GFP
200 µm


B

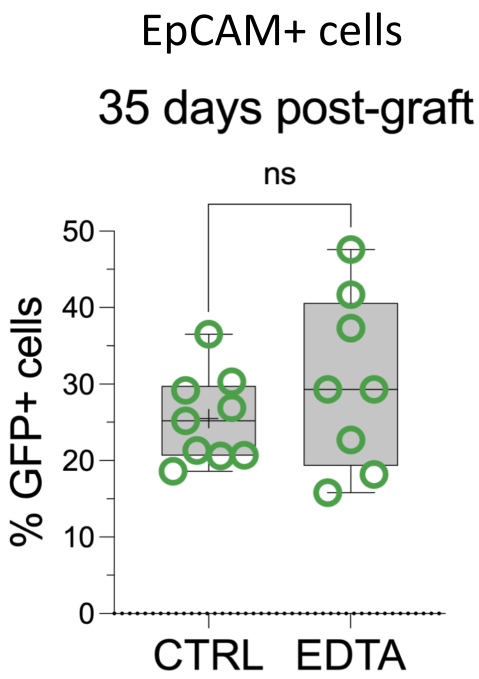

EpCAM+ cells
35 days post-graft
ns
% GFP+ cells
50
40
30
20
10
0
CTRL
EDTA

Table S1: Solution used as conditioning regimens prior engraftment experiments.

| | | Concentration | Exposition time (min) | Reference |
|---|---|---|---|---|
| **Trypsin** | | 0,25% | 15 | 25200056 |
| **Pronase** | | 1mg/mL | 30 | 10165921001 |
| **Dispase** | | 1U/mL | 30 | d4693 |
| **MMP9** | | 2% | 60 | ab285989 |
| **KLKs + Calpain-1** | KLK5 | 1-2,5ug/mL | 60 | 15950649 |
| | KLK13 | 2,5-5ug/mL | | 15955018 |
| | Calpain-1 | 0,1mg/mL | | C6108 |
| **EDTA** | | 0,2g/L | 30 | 15040066 |
| **CHAPS** | | 2,5mM | 15 | C3023-5G |
| **Polidocanol** | | 0,1% - 1% | 15 | Aetoxisclerol |

Table S2: Antibodies used for flow cytometry and immunofluorescence analysis.

| Target | Cell Type | Application | Reference | Dilution |
|---|---|---|---|---|
| Nkx2.1 (TTF1) | Lung progentors (vAFE) | FC (indirect) | ab76013 | 1:1000 |
| EPCAM | Epithelal cells | FC (direct) | 130-111-116 | 1:200 |
| CDH1 | Epithelal cells | IF | ab40772 | 1:500 |
| KRT5 | Basal cells | IF | ab64081 | 1:200 |
| Tubulin IV | Cilliated cells | IF | T7941 | 1:200 |
| MUC5AC | Goblet cells | IF | ab3649 | 1:200 |
| GFP | GFP positive cells | FC (indirect) / IF | ab6673 | 1:500 / 1:250 |
| *Secondary* | Anti-goat | FC (indirect) / IF | A32814 | 1:3000 / 1:1000 |
| | Anti-rabbit | | A32795 | |
| | Anti-mouse | | A32773 | |
| DAPI | Nucleus | IF | D9542 | 1:2500 |

Table S3: Primer pairs used for RTqPCR.

| Gene | Forward / Reverse | Tm |
|---|---|---|
| Nkx2.1 | CGGCATGAACATGAGCGGCAT | 64,27 |
| | GCCGACAGGTACTTCTGTTGCTTG | 64,12 |
| KRT5 | GGAGTTGGACCAGTCAACATC | 55,5 |
| | TGGAGTAGTAGCTTCCACTGC | 56,2 |
| MUC5AC | CATCTGCCAGCTGATTCTGA | 54,8 |
| | AAGACGCAGCCCTCATAGAA | 56 |
| CC10 | CATGAAACTCGCTGTCACCC | 56,3 |
| | GATGACACGCTGAAAGCTCG | 56,4 |
| FoxJ1 | GAGACAGGTTGTGGCGGATTGA | 59,5 |
| | ACTCGTATGCCACGCTCATCTG | 59,2 |
| PAX6 | GCGGAGTTATGATACCTACACC | 58,09 |
| | GAAATGAGTCCTGTTGAAGTGG | 57,24 |
| PAX8 | TCAACCTCCCTATGGACAGCTG | 61,48 |
| | GAGCCCATTGATGGAGTAGGTG | 60,49 |
| HNF4A | GCAGGCTCAAGAAATGCTTC | 57,73 |
| | GGCTGCTGTCCTCATAGCTT | 59,82 |
| SOX2 | GGCCATTAACGGCACACTGCC | 64,74 |
| | TTACTCTCCTCTTTTGCACCCCTCC | 64,03 |
| GAPDH | CTCTGCTCCTCCTGTTCGAC | 61,4 |
| | ACGACCAAATCCGTTGACTC | 57,3 |

**Video S1**

**Video S2**

**Video S3**
